# Phylodynamics of SARS-CoV-2 transmission in Spain

**DOI:** 10.1101/2020.04.20.050039

**Authors:** Francisco Díez-Fuertes, María Iglesias-Caballero, Sara Monzón, Pilar Jiménez, Sarai Varona, Isabel Cuesta, Ángel Zaballos, Michael M Thomson, Mercedes Jiménez, Javier García Pérez, Francisco Pozo, Mayte Pérez-Olmeda, José Alcamí, Inmaculada Casas

## Abstract

**Objectives:** SARS-CoV-2 whole-genome analysis has identified three large clades spreading worldwide, designated G, V and S. This study aims to analyze the diffusion of SARS-CoV-2 in Spain/Europe.

**Methods:** Maximum likelihood phylogenetic and Bayesian phylodynamic analyses have been performed to estimate the most probable temporal and geographic origin of different phylogenetic clusters and the diffusion pathways of SARS-CoV-2.

**Results:** Phylogenetic analyses of the first 28 SARS-CoV-2 whole genome sequences obtained from patients in Spain revealed that most of them are distributed in G and S clades (13 sequences in each) with the remaining two sequences branching in the V clade. Eleven of the Spanish viruses of the S clade and six of the G clade grouped in two different monophyletic clusters (S-Spain and G-Spain, respectively), with the S-Spain cluster also comprising 8 sequences from 6 other countries from Europe and the Americas. The most recent common ancestor (MRCA) of the SARS-CoV-2 pandemic was estimated in the city of Wuhan, China, around November 24, 2019, with a 95% highest posterior density (HPD) interval from October 30-December 17, 2019. The origin of S-Spain and G-Spain clusters were estimated in Spain around February 14 and 18, 2020, respectively, with a possible ancestry of S-Spain in Shanghai.

**Conclusions:** Multiple SARS-CoV-2 introductions have been detected in Spain and at least two resulted in the emergence of locally transmitted clusters, with further dissemination of one of them to at least 6 other countries. These results highlight the extraordinary potential of SARS-CoV-2 for rapid and widespread geographic dissemination.

## Introduction

The coronavirus disease 2019 (COVID-19) is an ongoing pandemic caused by severe acute respiratory syndrome coronavirus 2 (SARS-CoV-2) [1]. The symptom onset of the first case of COVID-19 was reported in December 1, 2019 in the city of Wuhan (China) and the initial outbreak was related to the Huanan seafood market [2-4]. SARS-CoV-2 infections have been reported in all the countries of Europe, causing from hundreds to thousands deaths in almost all them [5]. According to GISAID database, three larger clades of SARS-CoV-2 have been identified so far, based on marker variants [6]. Thus, S clade is characterized by the presence of L84S in ORF8 and mostly comprises sequences from North America. V clade is characterized by G251V in ORF3 and mostly comprises Asian and European sequences. Finally, G clade is characterized by D614G in S protein and largely comprises sequences from Europe. Other sequences have not been included in any of these clades because they lack the mentioned markers [6]. The situation in Europe seems to be dominated by the three clades, although G clade is the largest one in GISAID so far. More than 5000 whole genome sequences of viruses collected in Europe have been already sequenced with high coverage, allowing the performance of phylogenetic and evolutionary analyses to better understand the transmission dynamics within the most affected countries in Europe.

According to the European Centre for Disease Prevention and Control (ECDC), the first cases of SARS-CoV-2 infections in Europe were reported in the week of 26 January-1 February 2020, with 5 cases in France (one locally transmitted), 4 cases in Germany (all locally acquired), two imported cases in Italy, two imported cases in UK and one imported case in Finland [7]. In Spain, the first two cases were reported during the following week (2-8 February), the first one in La Gomera (Canary Islands) related to a known cluster in Germany and another one in Mallorca (Balearic Islands) from a British citizen who was initially in contact with a confirmed case in France returning from Singapore [8, 9]. The next 5 reported cases in Spain had all a travel history to Italy and were detected in Tenerife (Canary Islands), Catalonia, Castellon (Valencian Community) and Madrid before 26 February [10]. The following two days, Spain reported 18 new cases in Canary Islands, Catalonia, Andalusia and the Valencian Community and in the week of 1-7 March 2020, a total of 261 confirmed cases were reported in all the country [11]. The increasing number of whole genome sequences of SARS-CoV-2 circulating in Europe allows the development of phylodynamic studies to better understand these transmission dynamics of the virus in Spain and Europe and ideally to evaluate the proper functioning of surveillance systems.

## Methods

### Datasets

In order to investigate the transmission dynamics of SARS-CoV-2 in the most affected European countries, different datasets were retrieved from GISAID [6]. First, all the whole genome sequences (>29,000 bp) of the virus with high coverage were obtained (n=1590) to evaluate how the samples from Spain are distributed. Then, another dataset was analyzed including only whole genome sequences from Europe and from Wuhan, the city of the outbreak onset, to study the viral transmission dynamics within Europe. The last dataset was retrieved with whole genome sequences from Spain and from the countries where the first positives cases were reported, i.e. France, Germany, Italy and Finland. Sequences were aligned using MAFFT software and sequences were manually edited using AliView v1.26 [12, 13].

### Phylogenetic and evolutionary analyses

Phylogenies of large alignments were inferred by FastTree software v2.1.11 [14]. Root-to-tip genetic distances against sample collection dates were measured with TempEst v1.5.1 and Bayesian time-scaled phylogenetic analyses were performed with BEAST v1.10.4 to estimate the date and location of the most recent common ancestors (MRCA) as well as to estimate the rate of evolution of the virus [15, 16]. BEAST priors were introduced with BEAUTI v1.10.4 including an uncorrelated relaxed molecular clock model with a lognormal rate distribution and the exponential growth coalescent model of population size and growth [16]. Duplicated sequences were discarded from all the datasets. Markov chain Monte Carlo (MCMC) runs of 100 million states sampling every 10,000 steps were computed. The convergence of MCMC chains was checked using Tracer v.1.7.1, ensuring that the effective sample size (ESS) values were greater than 200 for each parameter estimated [17]. The maximum clade credibility (MCC) trees were obtained from the tree posterior distribution using TreeAnnotator after 10% burn-in [16].

### Viral transmission dynamics

Discrete trait evolutionary histories associated with phylogenies were analyzed with SpreaD3, generating klm files for visualizations in virtual globe software like Google Earth [18]. Well-supported rates between locations in the phylogeographic reconstructions were identified through a Bayes Factor by using the Bayesian stochastic search variable selection (BSSVS) procedure implemented in BEAST [19].

## Results

### Phylogeny of SARS-CoV-2 samples from Spain

A total of 28 whole genome sequences have been obtained from samples from Spain, including 14 from Madrid (12 of them sequenced by the University Hospitals 12 de Octubre, La Paz and Ramón y Cajal), five from the Valencian Community (sequenced by the FISABIO foundation), four from Castille and Leon, 2 from Castille-La Mancha and one each from Andalusia, Galicia and the Basque Country. These sequences branched in 3 different phylogenetic clusters (Figure 1A), 13 included in the G clade, two in the V clade and other 13 in the ancestral S clade (Figure 1B).

**Figure 1.**
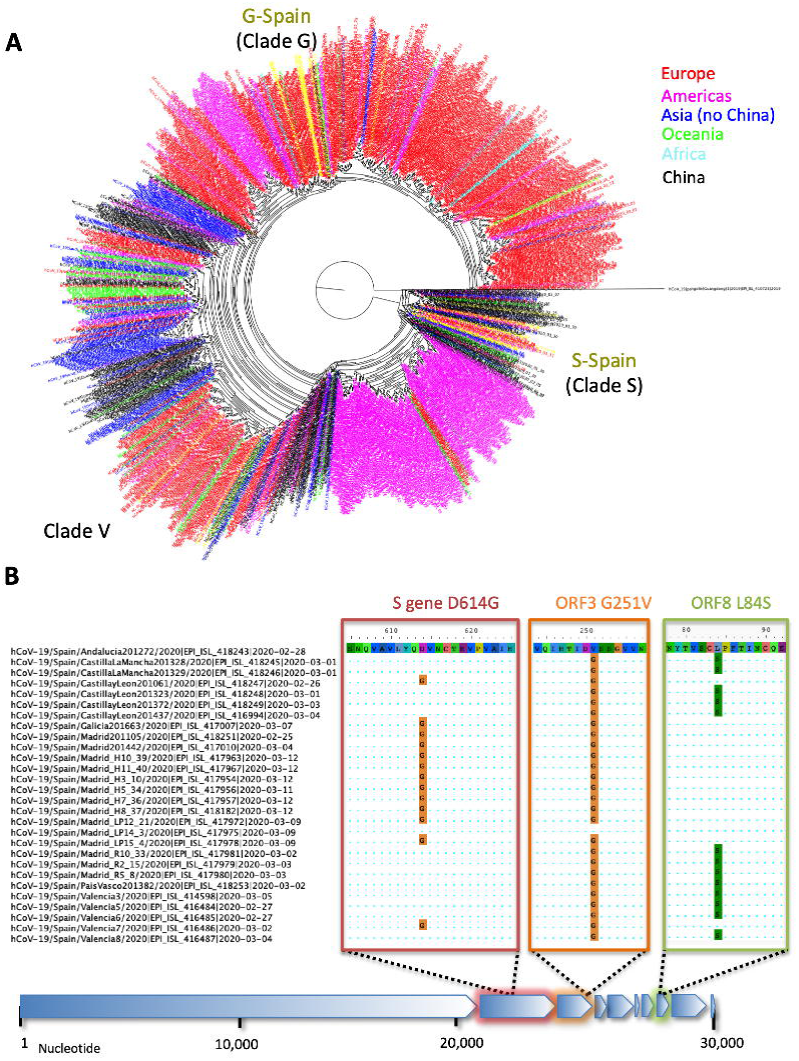
Phylogeny of the 1,590 whole genome sequences of SARS-CoV-2 with high coverage (>29,000 bp) and high quality available in GISAID [9] in March 20. The location of each sample is indicated by colors, differentiating sequences from Wuhan from the rest of sequences from Asia. Sequences from Spain are colored yellow in this figure to more easily distinguish them. FastTree2 software was used to build the phylogeny. The sequences EPI-ISL-410721 and -402131 of pangolin and bat coronaviruses were included to root the tree (A). Genetic characteristics on clade-defining markers for the 28 sequences from Spain available at the end of March. G clade contains D614G marker in the spike protein, V clade contains G251V in ORF3 and S clade contains L84S in ORF8 (B).

The phylogenies obtained for the dataset only comprising sequences from European countries revealed that the sequence EPI-ISL-406862, collected from a patient from Munich, Germany, in January 28, 2020, was basal to the G clade and at least six sequences from Spain, branched in the monophyletic G-Spain cluster (Figure 2A). In the case of the V clade, the sequences from Spain are integrated in a cluster where two sequences from Sweden (February 7, 2020) and France (January 23, 2020) were basal to the cluster formed by European sequences (Figure 2B). The rest of the sequences from Spain branched in the S clade and 11 of them were found in S-Spain cluster along with a basal sequence to this cluster from a virus collected in Shanghai on February 1, 2020 (Figure 2C).

**Figure 2.**
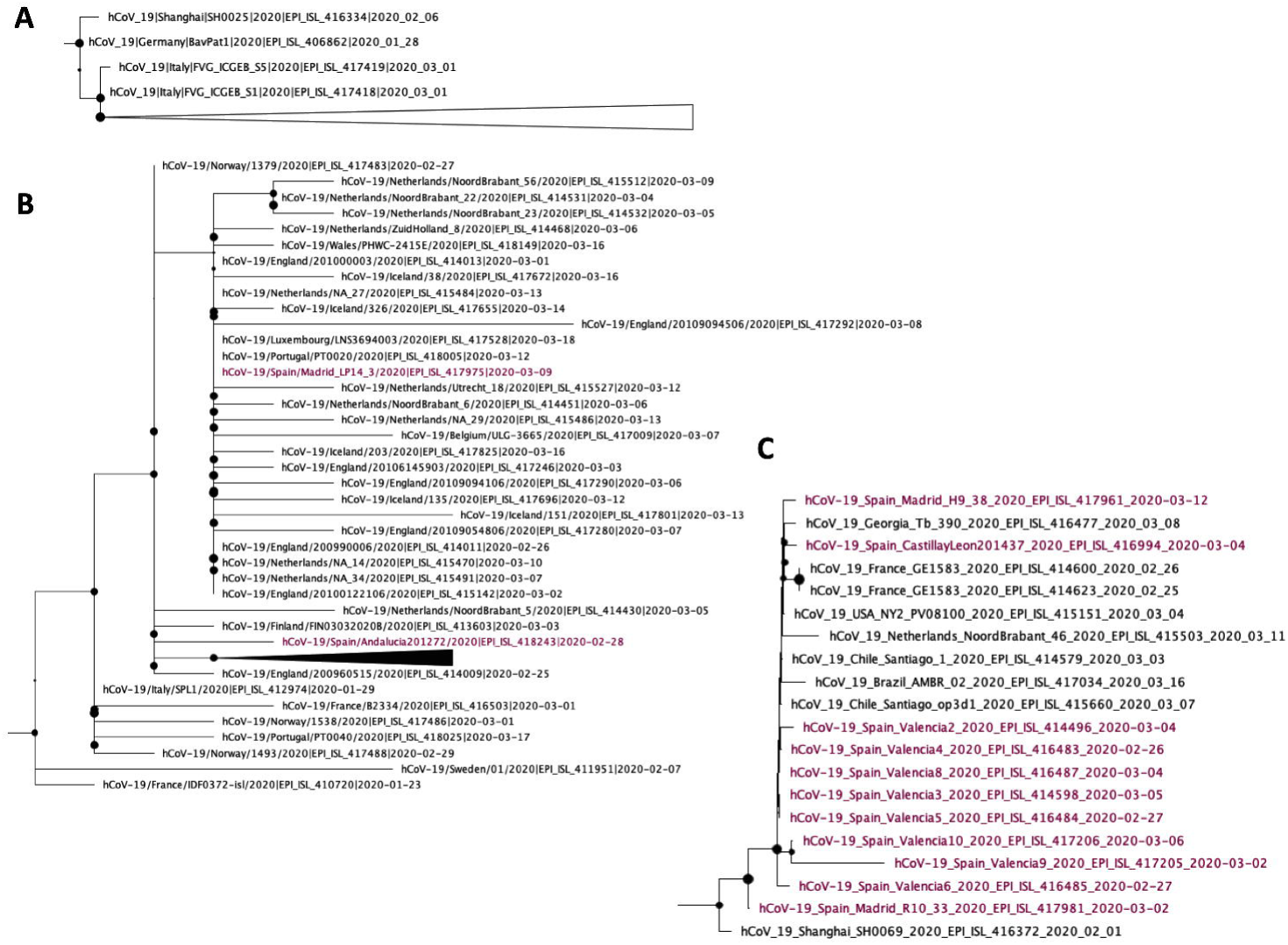
Detail of some sequences included in clades G (A), V (B) and S (C). FastTree2 software was used to build the phylogeny. Sequences from Spain were colored magenta. Some sequences of the trees were collapsed to facilitate the visualization of the origin of each cluster.

### Global expansion of SARS-CoV-2 worldwide

The MRCA estimated for the dataset including sequences from Wuhan and from the first European countries reporting cases was located in Wuhan (PP=0.99) around November 24, 2019 (95% highest posterior density (HPD) interval October 30-December 17, 2019). BF analysis revealed highly probable diffusions from Wuhan to all the European countries included in the dataset, i.e., England (BF>100; PP=1.00), Germany (BF>100; PP=1.00), France (BF>100; PP=1.00), Spain (BF>100; PP=1.00), Finland (BF>100; PP=0.99) and Italy (BF=29.1; PP=0.93). This dataset was also used to estimate the rate of evolution of SARS-CoV-2, which was 1.47×10^−3^ substitutions per site and per year (95% HPD interval 1.08×10^−3^ – 1.87×10^−3^). Results of the root-to-tip genetic distances against sample collection dates were measured for different phylogenetic clusters (Supplementary Figure 1).

### Origin of S and S-Spain clades

The most probable origin of the S phylogenetic cluster in Spain (Spain-S) was located in Shanghai (PP=0.65) around January 28, 2020 (95% HPD interval January 21-February 2, 2020). The MRCA of the S-Spain cluster, which also included 8 sequences from 6 other countries (USA, Netherlands, Chile, Brazil, Georgia and France) was located in Spain (PP=0.79) around February 14, 2020 (95% HPD interval February 4-23, 2020) (Figure 3; Supplement S1). The BF test indicated that the only probable diffusion between Asia and Europe was from China to the Netherlands (PP > 25; PP=0.87). In a different analysis where only sequences from Europe where included, the MRCA of this cluster was located in England (PP=0.95) and was dated around January 18, 2020 (95% HPD interval January 9-27).

**Figure 3.**
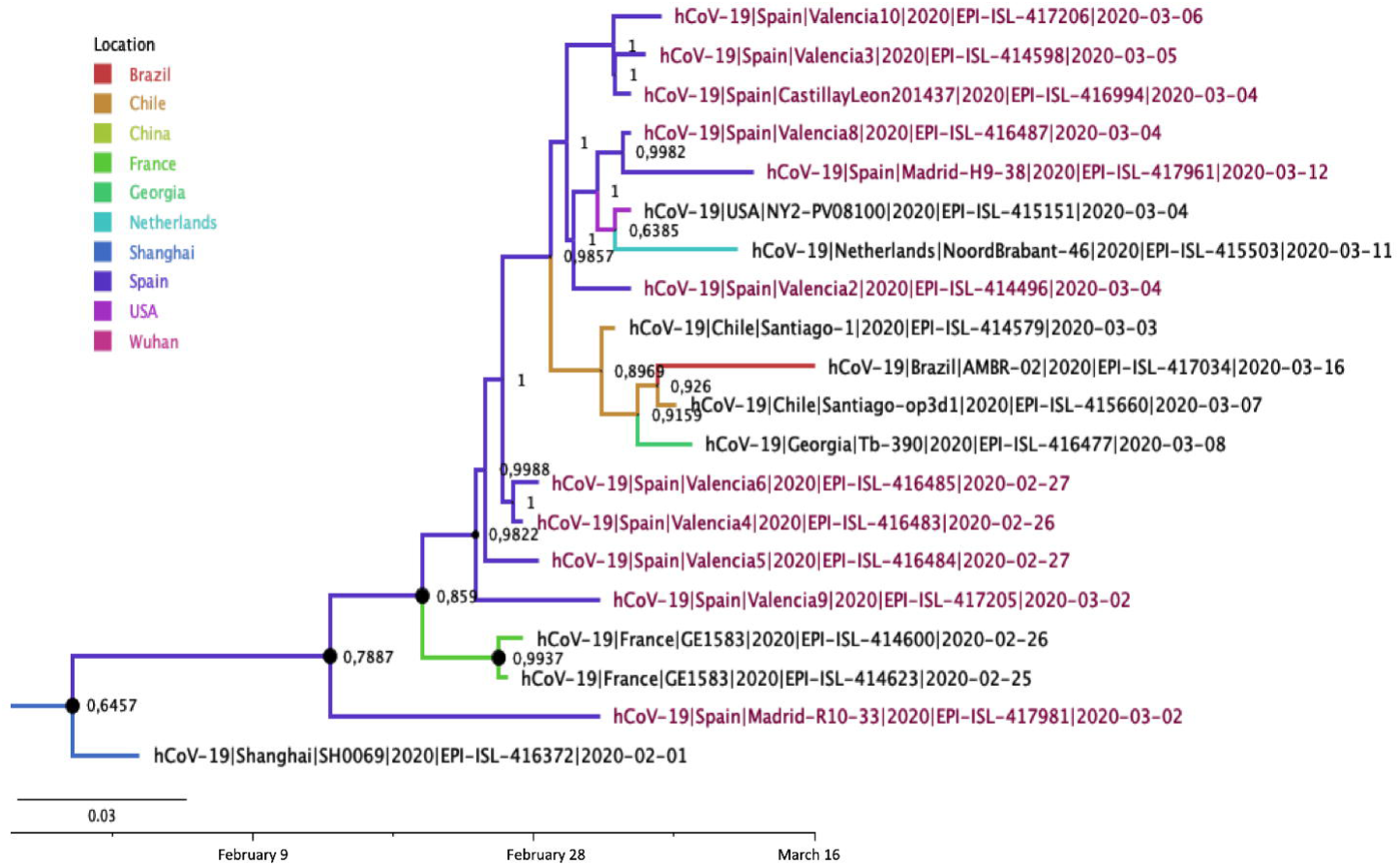
Maximum clade credibility (MCC) genealogies of S clade. Branch colors indicate the most probable location of the MRCA according to the legend and node labels indicate the posterior probability supporting the estimated MRCA location. Node support values are indicated by node size (only nodes with PP>0.9 are considered well-supported). Scale axis represents estimated dating of the MRCA for each cluster and label spacing defines exactly 9.16 days from the most recent sample included in the analysis (March 16, 2020, in this case).

### Origin of V clade

The origin of the V clade is uncertain, since no location reached values of PP>0.40. The tMRCA of the V clade in our analyses was around December 23, 2019 (95% HPD November 26, 2019 - January 11, 2020). The MCC of the V-clade revealed an earlier subcluster mainly transmitted in Asia and Australia and a more recent subcluster dominated by sequences from Europe (Figure 4).

**Figure 4.**
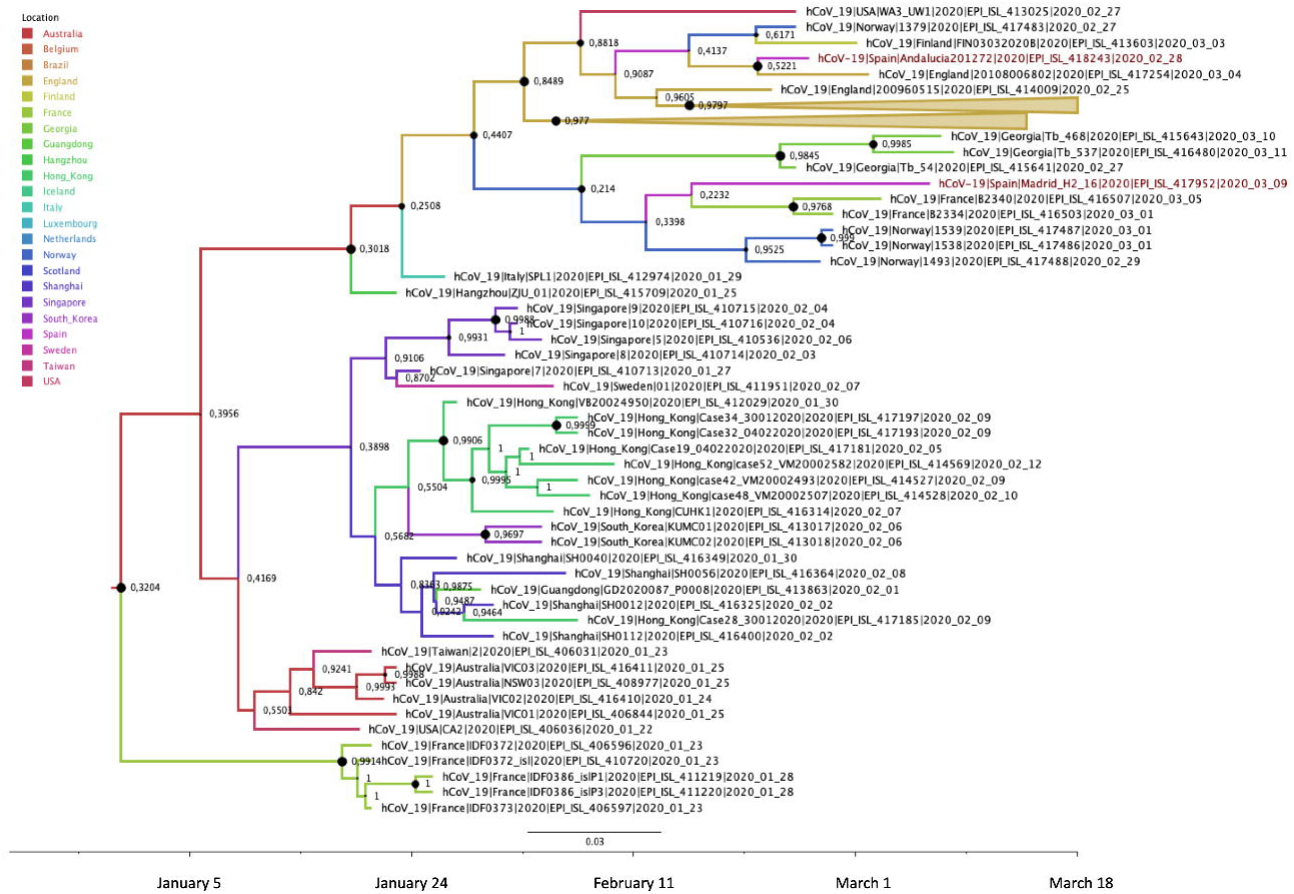
Maximum clade credibility (MCC) genealogies of V clade. Branch colors indicate the most probable location of the MRCA according to the legend and node labels indicate the posterior probability supporting the estimated MRCA location. Node support values are indicated by node size (only nodes with PP>0.9 are considered well-supported). Scale axis represents estimated dating of the MRCA for each cluster and label spacing defines exactly 9.16 days from the most recent sample included in the analysis (March 18, 2020, in this case).

Sequences from Spain were found in different subclusters of the V clade. Specifically, the sequences EPI-ISL-418243 and -417975 from Madrid and Andalusia, respectively, were found in a subcluster where the MCRA was located in England (PP=0.85) and dated around February 5, 2020 (95% HPD interval February 1-9, 2020). The other subcluster included the sequence EPI-ISL-417952 from Madrid, although the location of the MRCA of this group is uncertain between Norway (PP=0.21), England (PP=0.20) and Spain (PP=0.15). The tMRCA of this subcluster was dated on February 10 (95% HPD interval February 1-20, 2020) (Figure 4; Supplement S2). The BF analysis revealed two important hubs of transmission for the V clade within Europe. The main one was England with probable diffusion to Netherlands (BF>100; PP=0.96), Italy (BF=36.46; PP=0.78) and Scotland (BF=21.76: PP=0.68). The other hub was Norway, with probable exchange routes to England (BF>100; PP=0.91), Sweden (BF=20.13; PP=0.66), Finland (BF=6.01; PP=0.37), Luxembourg (BF=7.38; PP=0.42) and Belgium (BF=3.00; PP=0.23).

### Origin of G and G-Spain clades

The MRCA of G clade in Europe was located in England (PP=0.93) and dated on January 20, 2020 (95% HPD interval January 10-29, 2020). The sequence from the Bavarian patient (EPI-ISL-406862) was found at the root of the MCC, confirming the results obtained by the phylogenetic analysis. This clade includes approximately 50% of the whole genomes sequenced so far and the sequences from Spain are distributed throughout the clade rather than being clustered all together, indicating that several transmissions have been produced (Figure 6; Supplement S3). The MRCA of G-Spain cluster comprising the sequences EPI-ISL-418251, -417963, -417954, -417978, - 417956, -417967 from Spain was located in Spain (PP=0.95) and was dated on February 18, 2020 (95% HPD interval February 9-25, 2020) (Figure 5).

**Figure 5.**
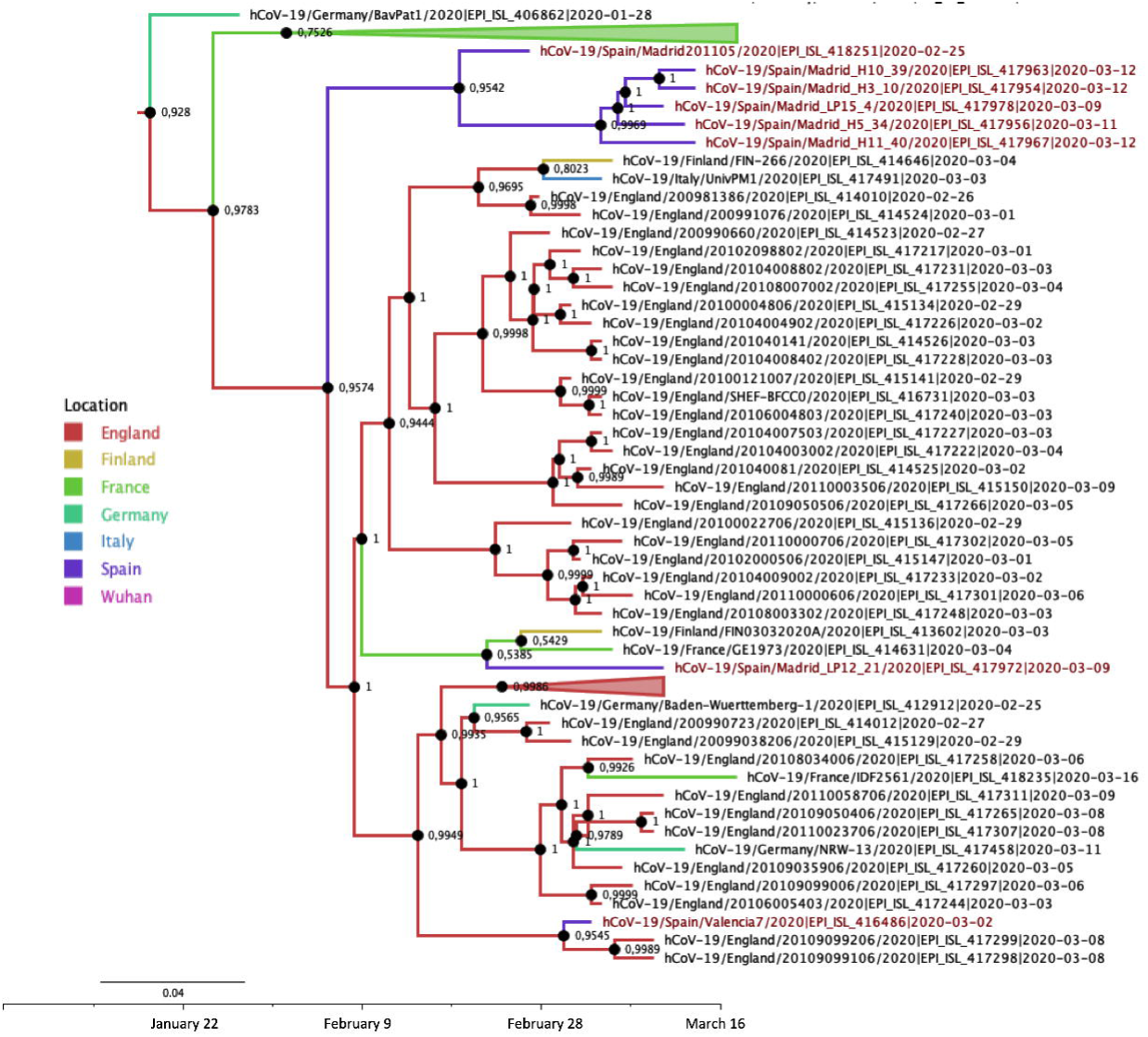
Maximum clade credibility (MCC) genealogies of G clade. Branch colors indicate the most probable location of the MRCA according to the legend and node labels indicate the posterior probability supporting the estimated MRCA location. Node support values are indicated by node size (only nodes with PP>0.9 are considered well-supported). Scale axis represents estimated dating of the MRCA for each cluster and label spacing defines exactly 9.16 days from the most recent sample included in the analysis (March 16, 2020, in this case).

## Discussion

The global analysis on the origin of the ongoing pandemic of SARS-CoV-2 indicated that the tMRCA was around November 24, 2019, with a 95% HPD interval from October 30 to December 17, 2019. This result is compatible with the epidemiologic information about the first case of COVID-19 reported in Wuhan, which dated the symptom onset in this patient on December 1, 2019 [3]. The estimated rate of evolution for SARS-CoV-2 was in the range of 1.08×10^−3^ – 1.87×10^−3^ substitutions per site per year, which is also comparable with the rates estimated for other epidemic coronaviruses, i.e. SARS-CoV (0.80×10^−3^ – 2.38×10^−3^) and MERS-CoV (0.88×10^−3^ – 1.37×10^−3^) [20, 21].

Taking into account the outbreak onset in Wuhan, it is worth pointing out that the city of Wuhan had direct flight connections with different European cities, including Paris (six weekly flights), London (three weekly flights) and Rome (three weekly flights) [22]. Regarding the V clade, five sequences from France are among the first representatives of this clade in Europe and were collected on January 23-28, 2020, from at least a couple of Chinese tourists from Wuhan [23]. In the present study two subclusters containing sequences from Spain were identified and the tMRCA was dated at the beginning of February and the MRCA of at least one of them was located in England. However, there is no scientific evidence supporting the association of the introduction of this clade in Spain to any known transmission cluster in Europe.

The origin of G clade in Europe has been associated with a known cluster of transmission in the state of Bavaria, Germany [24]. This outbreak began in January 19-22, 2020, with the contact of a “healthy German businessman” with a Shanghai resident who tested positive for SARS-CoV-2 on January 26 [9]. This cluster comprised at least 12 cases in Bavaria [24]. The present study dated the tMRCA of this clade on January 19, 2020, which is clearly compatible with the epidemiologic information. However, England was estimated as the origin of this cluster in spite of the fact that one representative sequence of the Bavarian cluster was included in the analysis (EPI-ISL-406962). This possible artifact could be explained by the low number of whole genome sequences available so far from Germany (n=8) compared with the sequences from England (n=58). A new cluster named G-Spain comprised of at least 6 sequences from Madrid, Spain, has been identified and the MRCA was located in Spain and dated around February 18, 2020.

The MRCA of the S phylogenetic cluster in Spain was located in Shanghai and dated around January 28. However, BF analyses did not support a direct transmission from Shanghai to Spain, suggesting that an ancestor of these viruses that are circulating in Spain was transmitted some time in the past in Shanghai but the introduction in Spain came from a different location. According to the same analysis, it is probable that the virus reached different European countries, including Spain, directly from Asia. The MRCA of the S-Spain clade was located in Spain and dated around February 14. This clade includes sequences from other 6 countries, suggesting its dissemination from Spain to USA, Netherlands, Chile, Brazil, Georgia and France. Epidemiologic investigations based on positive contact tracing have recently revealed that the first case detected in Europe was from a patient returning to France from Shanghai airport after a business trip in various cities, including Wuhan, in January 24 [23, 25].

A well-traced case transmitted in Europe is the cluster of the French skiing resort of Contamines-Monjoie. This cluster includes at least 13 infected individuals who are among the first cases reported in UK, France and Spain between 6 and 15 February, 2020 [9]. Recently, one of the sequences from this cluster has been released (EPI-ISL-410486) and its analysis revealed that it branches outside of G, V, and S clades. None of the sequences from Spain included in the present study has this topological feature, since all of them branch in one of the major clades. However, the second case reported in Spain (from Mallorca, Balearic Islands) is related to this cluster, so there is a possibility that a virus with these characteristics is also circulating in Spain but has not been sequenced yet [8, 9].

The main limitations of the present study are related to the fact that genomic sequences are being generated by diverse strategies following different steps that could affect the quality of the sequences. Different sample preparation techniques are being used, including overlapping amplicons, targeted capture where the viral RNA is enriched and metagenomic total RNA sequencing of rRNA depleted samples. The first two methods require less sequencing effort, but the possibility that some RNA molecules could be missed cannot be ruled out. On the contrary, the metagenomic approach is hypothesis-free, but implies a high number of sequencing reads. Another point to take into account is the sequencing strategies *per se*, since several approaches are being used, including Sanger sequencing and next generation sequencing platforms, such as iSeq, MiSeq, NextSeq and Novaseq from Illumina, MinION and GridION from Nanopore and IonTorrent from ThermoFisher [6]. All these technologies also have their own biases. Finally, the informatics employed to analyze the data is the step where more diversity of options are being identified. For all these reasons, some of the genetic differences observed between samples could be attributable to the error rate of sequencing platforms, indicating that genomes may be more similar than observed. On the other hand, the use of a reference genome to align the reads instead of following a *de novo* approach could mask some real genetic differences. In this sense, initiatives in *nf-core* are trying to provide best practice pipelines for the analysis of SARS-CoV-2 data in a peer-reviewed platform that includes some pipelines developed by the Bioinformatics Unit of the Instituto de Salud Carlos III [26-29]. Moreover, due to the public health threat caused by SARS-CoV-2, more genomic and epidemiologic data are generated every day and the results obtained in the present study should be seen as an early report about the current situation at the end of March 2020. However, studies like this about the dynamics of viral transmission are crucial to mitigate the pandemic because of phylodynamic analyses could help the public health authorities, identifying viral transmission routes and guiding the response measures.

## Acknowledgements

We thank Mar Molinero, Mónica González-Esguevillas, Sara Camarero, Diana Santos from the Respiratory and Influenza Unit for their exceptional technical support. We also thank the Rapid Response Unit from the National Center for Microbiology for their participation in the diagnosis of cases and the Microbiology Services at the following Hospitals for providing respiratory samples from patients: Hospital Universitario Virgen de las Nieves, Hospital General Universitario de Guadalajara, Hospital General de Segovia, Complejo Asistencial Universitario de Burgos, Hospital Santa María Nai, Hospital Universitario La Paz, Hospital Universitario Fundación Jiménez Díaz, Hospital Universitario 12 de Octubre, Hospital Universitario Ramón y Cajal, Hospital Txagorritxu de Vitoria, Hospital Clínico de Valencia and Consorcio Hospital General Universitario de Valencia. We gratefully acknowledge the authors, originating and submitting laboratories of the sequences from GISAID’s EpiCov™ Database on which this research is based. All submitters of data may be contacted directly via www.gisaid.org.

## Access to data

All the genomic sequences used in this manuscript have been deposited in GISAID and are fully accessible.

## Contribution

FDF made the phylogenetic studies and MIC, PJ, MJ, AZ obtained complete sequences from respiratory samples. SM, SV and ICu contributed with the Bioinformatic analysis of sequences. FDF, JGP and MPO wrote the first draft of the manuscript. MIC, ICu, AZ, MT, FP, JA and IC contributed to additional versions of the manuscript.

