## Supplementary figures and images for "Phylodynamics of SARS-CoV-2 transmission in Spain"

### SupplementaryFigureS1.png

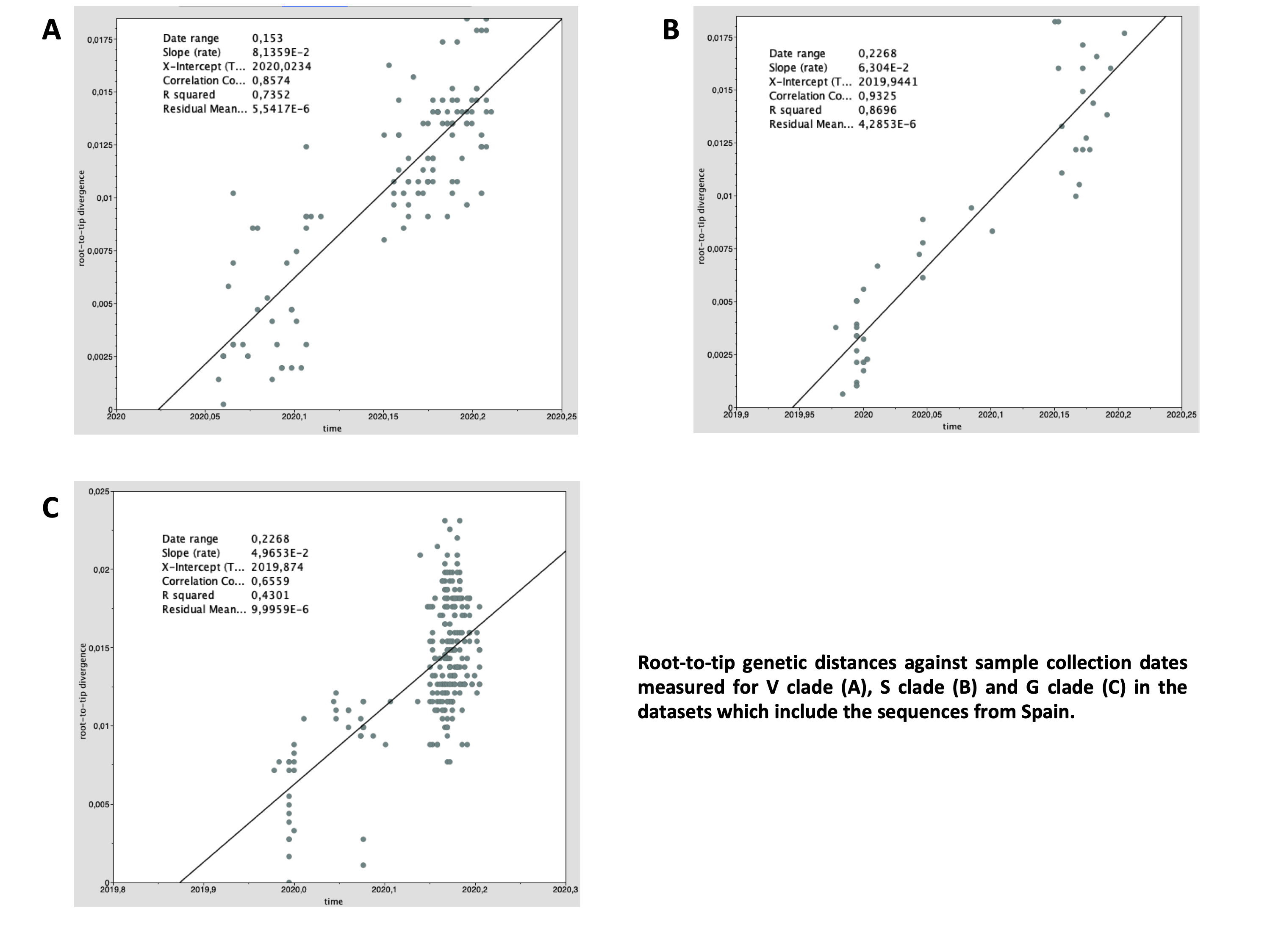
